# Lager yeast design through meiotic segregation of a fertile *Saccharomyces cerevisiae* x *Saccharomyces eubayanus* hybrid

**DOI:** 10.1101/2021.07.01.450509

**Authors:** Kristoffer Krogerus, Frederico Magalhães, Sandra Castillo, Gopal Peddinti, Virve Vidgren, Matteo De Chiara, Jia-Xing Yue, Gianni Liti, Brian Gibson

**Affiliations:** VTT Technical Research Centre of Finland, Tietotie 2, P.O. Box 1000, FI-02044 VTT, Espoo, Finland; Department of Biotechnology and Chemical Technology, Aalto University, School of Chemical Technology, Kemistintie 1, Aalto, P.O. Box 16100, FI-00076 Espoo, Finland; Institute for Research on Cancer and Ageing of Nice (IRCAN), CNRS UMR 7284, INSERM U1081, University of Nice Sophia Antipolis, Nice, France, 06107 Nice, France; Technische Universität Berlin, Chair of Brewing and Beverage Technology, Ackerstraße 76, 13355 Berlin, Germany; State Key Laboratory of Oncology in South China, Collaborative Innovation Center for Cancer Medicine, Sun Yat-sen University Cancer Center, Guangzhou, China

**Keywords:** Lager yeast, *S. eubayanus*, brewing, hybrid, tetraploid, sporulation

## Abstract

Yeasts in the lager brewing group are closely related and consequently do not exhibit significant genetic variability. Here, an artificial *Saccharomyces cerevisiae* × *Saccharomyces eubayanus* tetraploid interspecies hybrid was created by rare mating, and its ability to sporulate and produce viable gametes was exploited to generate phenotypic diversity. Four spore clones obtained from a single ascus were isolated, and their brewing-relevant phenotypes were assessed. These F1 spore clones were found to differ with respect to fermentation performance under lager brewing conditions (15 °C, 15 °Plato), production of volatile aroma compounds, flocculation potential and temperature tolerance. One spore clone, selected for its rapid fermentation and acetate ester production was sporulated to produce an F2 generation, again comprised of four spore clones from a single ascus. Again, phenotypic diversity was introduced. In two of these F2 clones, the fermentation performance was maintained and acetate ester production was improved relative to the F1 parent and the original hybrid strain. Strains also performed well in comparison to a commercial lager yeast strain. Spore clones varied in ploidy and chromosome copy numbers, and faster wort fermentation was observed in strains with a higher ploidy. An F2 spore clone was also subjected to 10 consecutive wort fermentations, and single cells were isolated from the resulting yeast slurry. These isolates also exhibited variable fermentation performance and chromosome copy numbers, highlighting the instability of polyploid interspecific hybrids. These results demonstrate the value of this natural approach to increase the phenotypic diversity of lager brewing yeast strains.

**Contribution to the field:** Lager beer fermentations have traditionally been carried out with natural *S. cerevisiae × S. eubayanus* hybrids. These strains possess both the ability to tolerate low temperatures and the ability to utilize efficiently wort sugars. However, being closely related, strains within the group exhibit limited phenotypic variability. Since the recent discovery of wild strains of *S. eubayanus*, it has been possible to generate lager yeast hybrids artificially, thereby increasing the genetic and phenotypic diversity of lager brewing strains. Here, to demonstrate the potential for further increased diversity, a constructed tetraploid hybrid was sporulated and spore clones derived from a single ascus were evaluated with respect to fermentation performance (sugar utilization, stress tolerance and volatile aroma synthesis). Meiosis introduced variability in a number of key parameters. One fertile spore clone from this F1 generation was sporulated to introduce further diversity and to demonstrate the potential of clone selection in steering phenotypes in a desirable direction. Genome instability of hybrids was observed, but this can be exploited to further increase diversity. This was demonstrated by assessing performance of variants isolated after ten consecutive rounds of fermentation. The approach allows for the introduction of phenotypic diversity without the need for targeted genetic modification.

## Introduction

Industrial lager yeast are derived from limited genetic stock. The *Saccharomyces pastorianus* yeast strains used for lager beer fermentation are natural interspecies hybrids of *S. cerevisiae* and *S. eubayanus* (Liti et al., 2005; Dunn and Sherlock, 2008; Nakao et al., 2009; Libkind et al., 2011; Walther et al., 2014; Gallone et al., 2019; Langdon et al., 2019). Exactly when or how the original hybridization occurred has been debated but the yeast in use today have originated from a limited number of strains which were isolated from lager fermentations in Central Europe in the late 19th century, when the use of pure cultures in brewing became common (Gibson and Liti, 2015; Gallone et al., 2019; Gorter De Vries et al., 2019). Lager strains originally arose after one or possibly two hybridization events that probably occurred when a domesticated strain of *S. cerevisiae* encountered a contaminant *S. eubayanus* strain during a traditional ale fermentation (Dunn and Sherlock, 2008; Walther et al., 2014; Baker et al., 2015; Monerawela et al., 2015; Okuno et al., 2015; Gallone et al., 2019; Salazar et al., 2019). A hybrid of the two species would have benefited by inheriting the superior fermentation performance of the ale strain, in particular the ability to use the key wort sugar maltotriose (Gibson et al., 2013), and the cryotolerance of the *S. eubayanus* strain (Gibson et al., 2013; Hebly et al., 2015). No naturally-occurring strains of *S. pastorianus* have been (knowingly) isolated since the 19^th^ century and it is unlikely that such strains will be found in the future. In addition, being interspecies hybrids and mostly aneuploid, existing strains exhibit low sporulation efficiency and spore viability. As such, increasing diversity through meiotic recombination and sexual mating, while possible, remains challenging (Gjermansen and Sigsgaard, 1981; Sanchez et al., 2012; Ota et al., 2018; Turgeon et al., 2021), in particular without the aid of targeted genetic intervention (Ogata et al., 2011; Xu et al., 2015; Alexander et al., 2016; Xie et al., 2018). Greater functional diversity amongst lager brewing yeast would be of advantage to the brewing industry, particularly as there now exists a demand for more efficient resource utilization and an increased trend for variety in beer characteristics (Kellershohn and Russell, 2015).

The discovery of *S. eubayanus* (Libkind et al., 2011) has, for the first time, allowed creation of *de novo S. cerevisiae* x *S. eubayanus* hybrids, and strains thus formed show strong fermentation performance compared to the parental strains as well as producing distinct flavour profiles (Hebly et al., 2015; Krogerus et al., 2015, 2016, 2017; Mertens et al., 2015; Alexander et al., 2016; Gorter de Vries et al., 2019). However, both sporulation efficiency and spore viability of *de novo* interspecies yeast hybrids are limited (Marinoni et al., 1999; Greig et al., 2002; Sebastiani et al., 2002; Bozdag et al., 2021) just as they are in the naturally occurring *S. pastorianus* strains. Post-zygotic infertility is a defining feature of allodiploid yeast (Naumov, 1996). However, sterility is not necessarily an obstacle to a hybrid’s fitness as clonal propagation allows such strains to survive indefinitely, and potentially to take advantage of the inherited phenotypes from both parental strains. The lager yeast *S. pastorianus* is, in fact, the classic example of this phenomenon (Kielland-Brandt and Nilsson-Tillgren, 1995). A number of factors may contribute to hybrid sterility, though recent research suggest that the inability of diverged chromosomes to undergo recombination is a key factor (Bozdag et al., 2021). Regardless of the mechanism involved, a consequence of sterility is that increased diversity through normal chromosomal recombination and cross-over during meiosis is not possible. However, there are mechanisms by which fertility can be recovered. One such route is endoreplication, whereby a sterile diploid hybrid doubles its genome content to become an allotetraploid capable of producing viable diploid spores (Sebastiani et al., 2002). The species barrier can similarly be overcome by mating diploid parents to generate an allotetraploid hybrid (Gunge and Nakatomi, 1972; Greig et al., 2002; Krogerus et al., 2017; Charron et al., 2019; Naseeb et al., 2021). Meiotic segregants derived from such crosses may be expected to vary considerably due to the segregation of orthologous genes from the parental strains and the creation of unique biochemical pathways and regulatory mechanisms (Landry et al., 2007), particularly if there exists a high degree of heterozygosity in the parental strains.

In an effort to produce diverse strains of *S. cerevisiae* x *S. eubayanus* for use in the brewing industry, a fertile tetraploid hybrid strain was here created through rare mating of an ale strain and the type strain of *S. eubayanus*. This hybrid strain was sporulated and four sibling spores derived from a single ascus were isolated. The brewing fermentation performance of each F1 meiotic segregant derived from this strain was characterized and compared with that of its siblings and the original tetraploid strain as well as the original diploid *S. cerevisiae* and *S. eubayanus* parents. Two of the F1 meiotic segregants were found to be fertile tetraploids and the isolation of F2 ascus siblings from the best-performing strain was carried out in order to further improve fermentation performance and flavour production. In an effort to assess the genotypic and phenotypic stability of the hybrids, one of the F2 spore clones was passaged 10 times in all-malt brewer’s wort and fermentation performance of this serial repitched yeast slurry and three single cell cultures derived from this population were assessed. Genome sequences were analysed to determine the main genetic changes (SNP, CNV, structural variation) associated with the observed changes. It is our contention that this approach is a feasible method for selectively producing natural, genetically and phenotypically diverse strains for the lager brewing industry.

## Materials & Methods

### Yeast strains

The two parental strains were *S. cerevisiae* VTT-A-81062 (VTT Culture Collection, Finland), an industrial brewer’s yeast strain, and the *S. eubayanus* type strain VTT-C12902 (VTT Culture Collection, Finland; deposited as CBS12357 at CBS-KNAW Fungal Biodiversity Centre). The industrial lager strain A-63015 was included to compare performance of *de novo* hybrids with that of an industrial strain. A tetraploid hybrid (A-81062 × C12902) strain was created in a previous study (Krogerus et al. 2017) and is deposited in the VTT Culture Collection as A-15225. Meiotic segregants of this strain derived from an individual ascus are deposited as A-15226, A-15227, A-15228 and A-15229. Further meiotic segregants of the tetraploid strain A-15227 are deposited as A-16232, A-16233, A-16234, A-16235. Strain A-16235 was further passaged through 10 consecutive batch fermentations in 15 °Plato wort, after which three single cell isolates were isolated from the yeast slurry. These isolates are here referred to as A235 G10 1-3.

### Generation of meiotic segregants

The meiotic segregants of the tetraploid interspecific hybrid A-15255 were obtained by first culturing A-15255 in YPM medium (1% yeast extract, 2% peptone, 4% maltose) at 20 °C overnight. It was then transferred to pre-sporulation medium (0.8% yeast extract, 0.3% peptone, 10% glucose) at a starting OD600 of 0.3 and allowed to grow for 20 hours at 20 °C. The yeast was then washed with 1% potassium acetate and a thick suspension was plated onto sporulation agar (1% potassium acetate and 2% agar). The yeast was allowed to sporulate for 7 days at 25 °C. Meiotic segregants were obtained by dissecting tetrad ascospores treated with Zymolyase 100T (US Biological, USA) on YPD agar with a micromanipulator. Spore viability was calculated based on the amount of colonies formed from the dissection of up to 20 tetrads.

### DNA content by flow cytometry

Flow cytometry was performed on the yeast strains essentially as described by Haase & Reed (2002) and Krogerus et al. (2016). Briefly, the yeast strains were grown overnight in YPD medium (1% yeast extract, 2% peptone and 2% glucose), after which cells were fixed in 70% ethanol, treated with RNAse A (0.25 mg mL^-1^) and Proteinase K (1 mg mL^-1^), stained with SYTOX Green (2 μM; Life Technologies, USA), and their DNA content was determined using a FACSAria cytometer (Becton Dickinson). Measurements were performed on duplicate independent yeast cultures, and 100 000 events were collected per sample during flow cytometry.

### Genome sequencing and analysis

Genome assemblies of both parent strains, *S. cerevisiae* A-81062 and *S. eubayanus* C-12902, were first obtained in order to create a reference genome to which sequencing reads from the hybrid strains could be aligned. A long-read assembly of *S. eubayanus* C-12902 was obtained from Brickwedde et al. (2018). *S. cerevisiae* A-81062 has been sequenced previously by our group using an Oxford Nanopore Technologies MinION (Krogerus et al., 2019) and with Illumina technology (Krogerus et al., 2016). Reads were accessed from SRR9129759 and SRR2911435, respectively. Here, the long reads were *de novo* assembled using the LRSDAY (version 1.4) pipeline (Yue and Liti, 2018). The initial assemblies were produced with smartdenovo (available from https://github.com/ruanjue/smartdenovo) using default settings. The assembly was first polished with medaka (1.2.0; available from https://github.com/nanoporetech/medaka), followed by two rounds of short-read polishing with Pilon (version 1.23; Walker et al., 2014). Alignment of long reads for medaka was performed with minimap2 (version 2.17-r941; Li, 2018). The contigs in the polished assemblies were then scaffolded with Ragout (version 2.3; Kolmogorov et al., 2014) to *S. cerevisiae* S288C (R64-2-1). Because of the relatively high heterozygosity of *S. cerevisiae* A-81062, two haplotypes were further produced through phasing in WhatsHap (version 1.0; Martin et al., 2016). Short reads were first mapped to above scaffolds, and variants were called with FreeBayes (version 1.32; Garrison and Marth, 2012). Long reads were also mapped to the above scaffolds with minimap2, and the resulting VCF and long-read BAM files were then passed to WhatsHap. The two haplotypes of *S. cerevisiae* A-81062 were then extracted from the resulting phased VCF. Assembly statistics are available in Supplementary Table S1 and Supplementary Figure S1, while the A-81062 assembly is available as Supplementary Data 1. A reference genome for the analysis of the hybrid strains was produced by concatenating *S. cerevisiae* A-81062 haplotype 1 with the obtained assembly of *S. eubayanus* C-12902. The genomes of both parent strains were also separately annotated using MAKER2 (Holt and Yandell, 2011) as implemented in the LRSDAY pipeline. A horizontal gene transfer event from *Torulaspora microellipsoides* in the *S. cerevisiae* A-81062 genome was identified by mapping chromosome XV to scaffold FYBL01000004.1 of *T. microellipsoides* CLIB830 (NCBI GCA_900186055.1; Galeote et al., 2018) using minimap2 (with ‘-x asm20’ parameter). Alignments were visualized with the ‘pafr’-package for R (https://github.com/dwinter/pafr).

The tetraploid hybrid A-15225 and all derived spore clones and G10 isolates were sequenced by Biomedicum Genomics (Helsinki, Finland). The sequencing of A-15225 has been described previously in Krogerus et al. (2017) and reads are available from NCBI-SRA SRR5141258 (referred to as ‘Hybrid H1’). In brief, an Illumina KAPA paired-end 150 bp library was prepared for each strain and sequencing was carried out with a NextSeq 500 instrument. The newly described Illumina sequencing reads have been submitted to NCBI-SRA under BioProject number PRJNA357993. Paired-end reads from the NextSeq 500 sequencing were trimmed and filtered with fastp using default settings (version 0.20.1; Chen et al., 2018). Trimmed reads were aligned to the concatenated reference genome described above using BWA-MEM (Li and Durbin, 2009), and alignments were sorted and duplicates were marked with sambamba (version 0.7.1; Tarasov et al., 2015). Variants were jointly called in the twelve hybrid strains using FreeBayes (version 1.3.2; Garrison and Marth, 2012). Variant calling used the following settings: --min-base-quality 30 --min-mapping-quality 30 --min-alternate-fraction 0.25 --min-repeat-entropy 0.5 --use-best-n-alleles 70 -p 2. The resulting VCF file was filtered to remove variants with a quality score less than 1000 and with a sequencing depth below 10 per sample using BCFtools (Li, 2011). The haplotype blocks in the meiotic segregants were obtained from the filtered VCF file by clustering consecutive reference (haplotype 1) or alternative (haplotype 2) allele calls using the vcf_process.pl script from https://github.com/wl13/BioScripts. Variants were annotated with SnpEff (version 4.5covid19; Cingolani et al., 2012). Visualizations were performed in R using the ‘karyoploter’ package (Gel and Serra, 2017). Chromosome copy numbers were estimated based on the median coverage in 10kb windows across each contig in the reference genome as calculated with mosdepth (version 0.2.6; Pedersen and Quinlan, 2018). Alignment of reads to the right arm of *S. cerevisiae* chromosome XV was visualized with samplot (https://github.com/ryanlayer/samplot).

Structural variations in the *S. cerevisiae* A-81062 parent strain were identified using long sequencing reads. Long reads were first aligned to the *de novo* assembly produced above using NGMLR (version 0.2.7; Sedlazeck et al., 2018), after which structural variations were called from the alignment using Sniffles (version 1.0.12; Sedlazeck et al., 2018). Variants were annotated with SnpEff (Cingolani et al., 2012). Gene ontology enrichment analysis on the set of genes affected by heterozygous structural variants was carried out with YeastMine (Balakrishnan et al., 2012). Structural variations in the hybrid strains were estimated from split and discordant Illumina reads using LUMPY (Layer et al., 2014) and genotyped with svtyper (Chiang et al., 2015) as implemented in smoove (version 0.2.6; https://github.com/brentp/smoove). Variations in all twelve hybrid strains were jointly called, and the resulting VCF was filtered to remove sites with an imprecise breakpoint or a quality score less than 100 using BCFtools (Li, 2011).

### Fermentations

Yeast performance was determined in fermentations carried out at 15 °C in a 15 °Plato all-malt wort. Yeast was propagated essentially as described previously (Krogerus et al. 2015) with the use of a ‘Generation 0’ fermentation prior to the actual experimental fermentations. The experimental fermentations were carried out in duplicate, in 2-L cylindroconical stainless steel fermenting vessels, containing 1.5 L of wort medium. The 15 °Plato wort was produced at the VTT Pilot Brewery from barley malt and was oxygenated to 15 mg L^−1^ prior to pitching. Yeast was inoculated at a rate of 5g L^−1^ to the wort. Wort samples were drawn regularly from the fermentation vessels aseptically, and placed directly on ice, after which the yeast was separated from the fermenting wort by centrifugation (9000 × *g*, 10 min, 1 °C). Samples for yeast-derived flavour compounds and fermentable sugars were taken from the beer.

### Wort and beer analysis

The specific gravity, alcohol level (% v/v) and pH of samples was determined from the centrifuged and degassed fermentation samples using an Anton Paar Density Meter DMA 5000 M (Anton Paar GmbH, Austria) with Alcolyzer Beer ME and pH ME modules (Anton Paar GmbH, Austria). Concentrations of fermentable sugars (glucose, fructose, maltose and maltotriose) were measured by HPLC using a Waters 2695 Separation Module and Waters System Interphase Module liquid chromatograph coupled with a Waters 2414 differential refractometer (Waters Co., Milford, MA, USA). An Aminex HPX-87H Organic Acid Analysis Column (300 × 7.8 mm, Bio-Rad) was equilibrated with 5 mM H_2_SO_4_ (Titrisol, Merck, Germany) in water at 55 °C and samples were eluted with 5 mM H_2_SO_4_ in water at a 0.3 ml/min flow rate.

Yeast-derived flavour compounds were determined by headspace gas chromatography with flame ionization detector (HS-GC-FID) analysis. 4 mL samples were filtered (0.45 µm), incubated at 60 °C for 30 minutes and then 1 mL of gas phase was injected (split mode; 225 °C; split flow of 30 mL min^-1^) into a gas chromatograph equipped with a FID detector and headspace autosampler (Agilent 7890 Series; Palo Alto, CA, USA). Analytes were separated on a HP-5 capillary column (50m × 320 µm × 1.05 µm column, Agilent, USA). The carrier gas was helium (constant flow of 1.4 mL min^-1^). The temperature program involved 50 °C for 3 min, 10 °C min^-1^ to 100 °C, 5 °C min-1 to 140 °C, 15 °C min^-1^ to 260 °C and then isothermal for 1 min. Compounds were identified by comparison with authentic standards and were quantified using standard curves. 1-Butanol was used as internal standard.

### Yeast analysis

The yeast dry mass content of the samples (i.e. yeast in suspension) was determined by washing the yeast pellets gained from centrifugation with 25 mL deionized H_2_O and then suspending the washed yeast in a total of 6 mL deionized H_2_O. The suspension was then transferred to a pre-weighed porcelain crucible, and was dried overnight at 105° C and allowed to cool in a desiccator before the change of mass was measured. Yeast viability was measured from the yeast that was collected at the end of the fermentations using a Nucleocounter® YC-100^™^ (ChemoMetec). Flocculation of the yeast strains was evaluated using a modified Helm’s assay (D’Hautcourt and Smart, 1999).

### Data and statistical analysis

Data and statistical analysis on the fermentation and yeast data was performed with R (http://www.r-project.org/). One-way ANOVA and Tukey’s post hoc test was performed using the ‘agricolae’ package (De et al., 2017). Values were considered significantly different at p < 0.05. Heatmaps were drawn with the ‘pheatmap’ package (Kolde, 2015).

## Results

### Hybrid generation and genomic analysis

The set of 12 *de novo* hybrid strains used in this study were generated according to Figure 1. The tetraploid interspecies hybrid A225, from a cross between the *S. cerevisiae* A62 ale strain and the *S. eubayanus* C902 type strain, was obtained with ‘rare mating’ in a previous study (Krogerus et al., 2017). This interspecies hybrid sporulated efficiently and spores showed a viability of 55%. A set of four F1 segregants (A226-A229), all derived from the same ascus, were isolated. F1 segregant A227 also sporulated efficiently, and a set of four F2 segregants (A232-A235) were derived from this strain. F2 segregant A235 was further subjected to ten consecutive batch fermentations in 15 °P wort (corresponding to approximately 30-40 cells doublings), and three single cell isolates (A235 G10 1-3) were randomly selected from the resulting yeast population.

**Figure 1.**
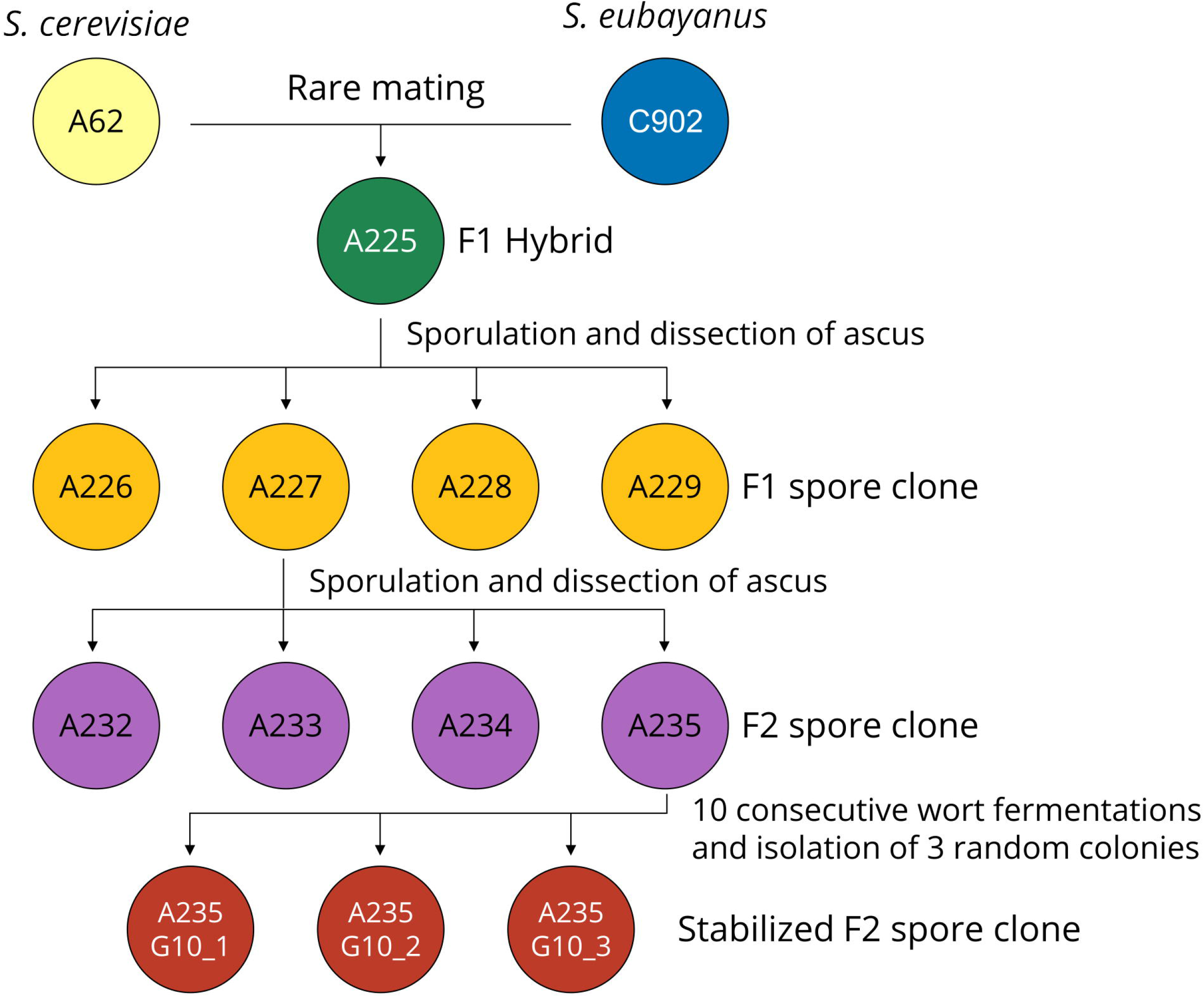
Overview of the yeast strains generated in this study.

For the genomic analysis of the hybrid strains, a new *de novo* assembly of parent strain *S. cerevisiae* A62 was produced for use as reference genome. The genome of A62 has been assembled previously using a hybrid strategy (assembly from 150 bp Illumina reads, and scaffolding with PacBio reads) (Krogerus et al., 2016). Here, a long-read assembly was instead produced with smartdenovo using reads generated with the Oxford Nanopore MinION from our previous study (Krogerus et al., 2019). The assembly was polished once with long reads in Medaka, and twice with Illumina reads in Pilon. The resulting assembly consisted of 21 scaffolds (including the 16 chromosomes and mitochondrial DNA) and spanned a genome size of 12.68 Mbp (assembly statistics available in Supplementary Table S1 and Supplementary Figure S1). A total of 29517 heterozygous single nucleotide polymorphisms were detected, corresponding to a heterozygosity of around 0.23%. The heterozygous SNPs were phased in whatshap using the long sequencing reads, and the two haplotypes were extracted. 90% of the heterozygous SNPs (26569) were phased into a total of 29 blocks (1.45 per scaffold). The first haplotype was selected to be used as reference for the *S. cerevisiae* A62 parent strain. The reference genome for the *S. eubayanus* C902 parent strain was obtained from Brickwedde et al. (2018). The genomes were separately annotated using the MAKER-based pipeline in LRSDAY, and a total of 5945 and 5430 protein-coding genes were detected, respectively. For analysis of the hybrid strains produced in this study, a concatenated reference genome of *S. cerevisiae* A62 and *S. eubayanus* C902 was used.

#### Chromosome copy number variation

Chromosome copy numbers of the F1 hybrid and derived spore clones were estimated based on median coverage of the sequencing reads and flow cytometry with SYTOX Green-staining (fluorescence histograms available in Supplementary Figure S2). Diversity in both ploidy and individual chromosome copy numbers were observed (Figure 2). The two parent strains have been previously shown to be diploid (Krogerus et al., 2016). The genome of the F1 hybrid A225 consisted of two copies of each chromosome from *S. cerevisiae* and *S. eubayanus*. An exception was the *S. cerevisiae* chromosome III with only one copy, likely related to the rare mating. The mitochondrial genome in A225 and derived strains was inherited from *S. eubayanus*.

**Figure 2.**
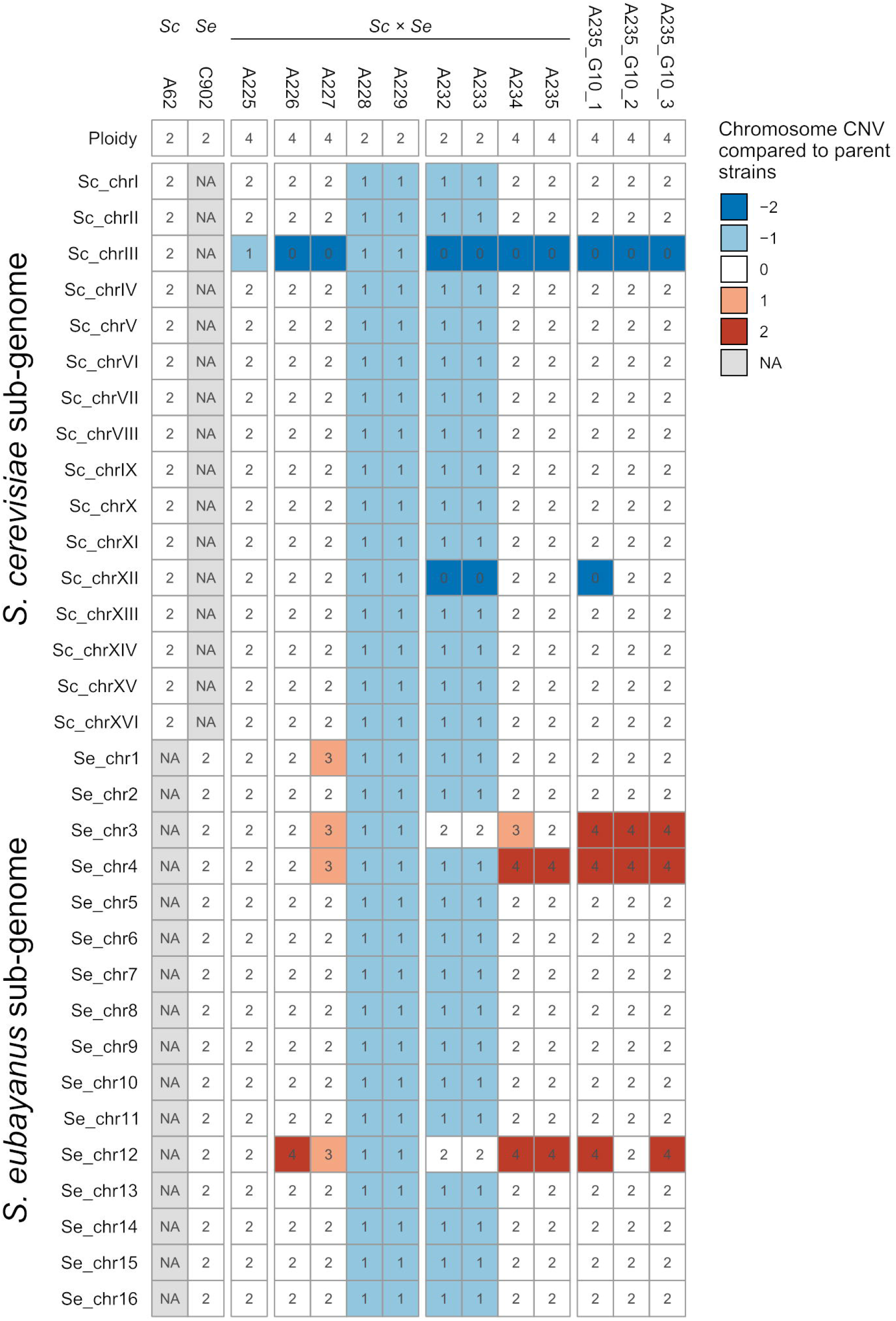
Chromosome copy numbers and ploidy of the parent and hybrid strains. Chromosome copy number variations (CNV) in the *S. cerevisiae* A-81062 (top) and *S. eubayanus* C12902 (bottom) sub-genomes of the hybrid strains compared to the parent strains (the numbers inside the cells indicate the estimated absolute chromosome copy number). A blue color indicates a chromosome loss, while a red color indicates a chromosome duplication compared to the parent strain (e.g., −1 corresponds to one less chromosome in the hybrid compared to the parent strain). NA, not available.

The four F1 hybrid spores were found to include two tetraploid strains (A226 and A227) and two diploid strains (A228 and A229). The diploid strains contained one copy of each chromosome from both *S. cerevisiae* and *S. eubayanus* (Figure 2). The tetraploid F1 strains contained two copies of each chromosome. Exceptions included chromosome I (three copies from *S. eubayanus* in strain A227), chromosome III (no copy from *S. cerevisiae* in A226 and A227, and an additional copy from *S. eubayanus* in A227), chromosome IV (with an additional copy from *S. eubayanus* in A227) and chromosome XII (four and three copies of the *S. eubayanus* form in A226 and A227, respectively).

Of the four F2 segregants derived from A227, two were again diploid (A232 and A233) and two were tetraploid (A234 and A235). The diploid strains contained one copy of each chromosome from *S. cerevisiae* and *S. eubayanus*, the exception being chromosome III for which only *S. eubayanus* was represented (2 copies) due to the lack of the corresponding *S. cerevisiae* chromosome in the parental A227 strain. Similarly, the diploid F2 hybrids did not contain the *S. cerevisiae* chromosome XII but this was compensated by having two copies of the *S. eubayanus* form of the chromosome. The tetraploid F2 hybrids possessed two copies of both the *S. cerevisiae* and *S. eubayanus* chromosomes with the exception that *S. cerevisiae* chromosome III was absent (three and two copies of the *S. eubayanus* form were present in A234 and A235 respectively). Both strains contained four copies of *S. eubayanus* chromosomes IV and XII from both parental species (Figure 2).

Further chromosome copy number variation was observed in the G10 isolates of A235, and interestingly all three single cell isolates exhibited different profiles (Figure 2). Compared to A235, all three single cell isolates carried an additional two copies of *S. eubayanus* chromosome III. Furthermore, A235 G10 1 had lost both copies of *S. cerevisiae* chromosome XII, while A235 G10 2 had lost two out of four copies of *S. eubayanus* chromosome XII.

#### Single nucleotide and structural variations

Recombination was observed within the parental sub-genomes of the F1 spore clones. As the reference genome of *S. cerevisiae* A62 was phased, recombination in the *S. cerevisiae* sub-genome of the F1 spore clones could be easily observed by presence of either of the two haplotype blocks (Figure 3). Such visualization could not be produced for the *S. eubayanus* sub-genome because of a considerably lower heterozygosity level (0.002%; Hebly et al., 2015). Of the 24726 heterozygous SNPs observed in the A225 F1 hybrid (24117 and 609 in the *S. cerevisiae* and *S. eubayanus* sub-genomes, respectively), 23017 segregated in a 2:2 pattern in the four F1 spore clones. Compared to A225, a total of 132 *de novo* SNPs were detected in the four F1 spore clones. Of these, 22 were missense mutations and two conservative in-frame insertions (Table 2). A 2:2 segregation pattern was observed for many of these SNPs (i.e. mutation present in two out of four spore clones), suggesting that the mutation might have been heterozygous in the F1 hybrid, despite showing a 0/0 genotype (i.e. only reference allele detected), and therefore not a true *de novo* mutation.

**Figure 3.**
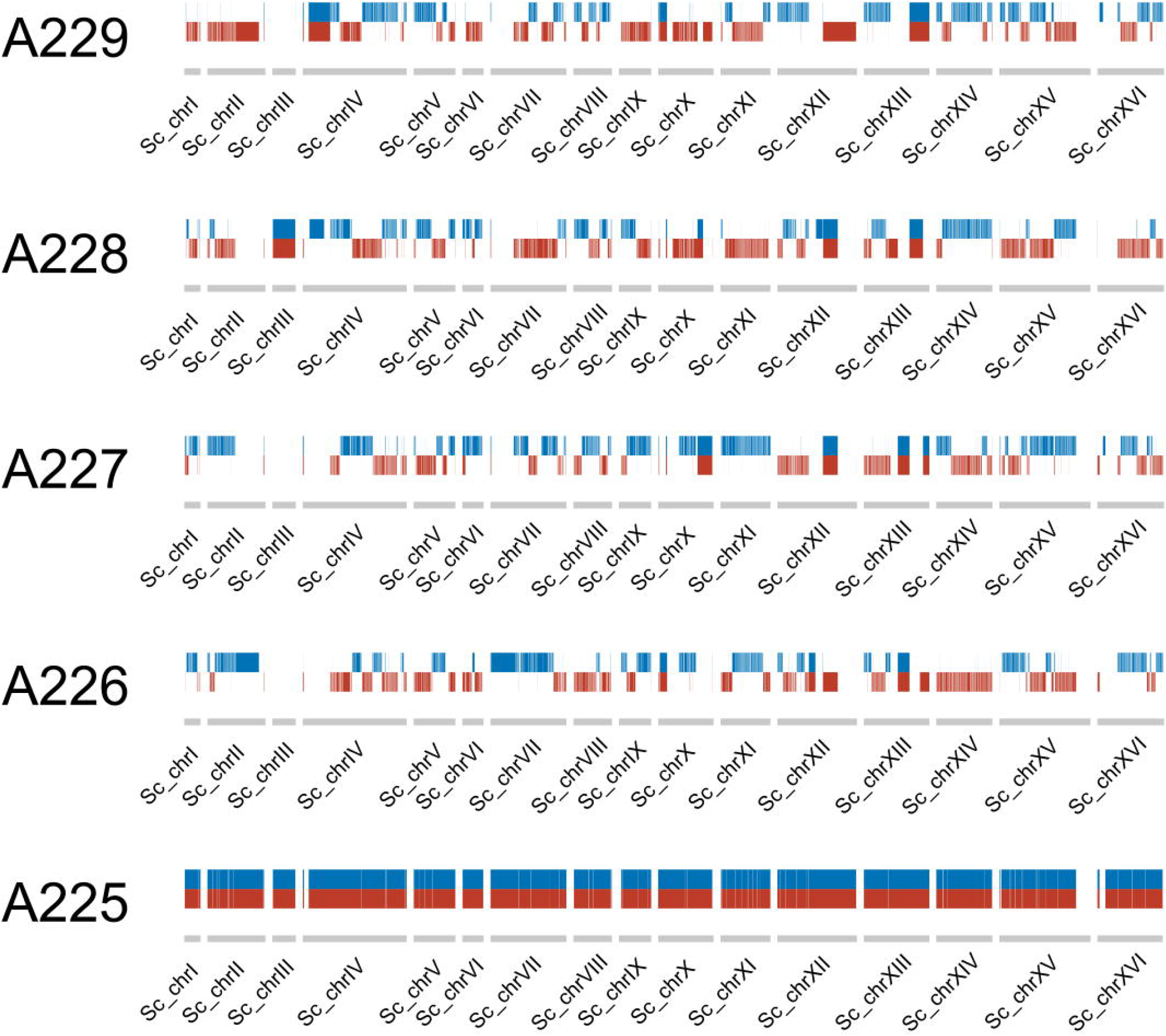
Haplotype blocks (red and blue) in the *S. cerevisiae* sub-genome of the F1 hybrid and the four F1 spore clones.

**Table 1.** Strains used in this study and their spore viabilities, flocculation potential, and post-fermentation viability. Spore viability was assessed by dissecting at least 16 tetrads by micromanipulation and observing colony formation after 4 days (YPM media, 24°C). ND: not determined. NA: not available.

| VTT Code | Short Code | Strain | Spore viability (%) | Flocculation potential (%) | Post-fermentation viability (%) |
| --- | --- | --- | --- | --- | --- |
| A-81062 | A62 | *S. cerevisiae* ale strain | 8 | 99 ± 0.0 | 97 ± 0.2 |
| A-63015 | A15 | *S. pastorianus* lager strain | 0 | ND | 92 ± 0.4 |
| C-12902 | C902 | *S. eubayanus* type strain | 96 | 3.0 ± 3.1 | 64 ± 2.0 |
| A-15225 | A225 | Hybrid of A-81062 and C-12902 | 55 | 92 ± 1.3 | 76 ± 2.0 |
| A-15226 | A226 | Meiotic segregant of A-15225 | 63 | 96 ± 1.1 | 71 ± 3.4 |
| A-15227 | A227 | Meiotic segregant of A-15225 | 95 | 4.2 ± 0.1 | 76 ± 0.5 |
| A-15228 | A228 | Meiotic segregant of A-15225 | 0 | 88 ± 0.8 | 98 ± 0.1 |
| A-15229 | A229 | Meiotic segregant of A-15225 | 0 | 2.8 ± 4.0 | 95 ± 0.1 |
| A-16232 | A232 | Meiotic segregant of A-15227 | 78 | 0.6 ± 0.1 | 94 ± 0.1 |
| A-16233 | A233 | Meiotic segregant of A-15227 | 0 | 1.0 ± 4.9 | 93 ± 0.2 |
| A-16234 | A234 | Meiotic segregant of A-15227 | 78 | 0.0 ± 3.1 | 17 ± 2.1 |
| A-16235 | A235 | Meiotic segregant of A-15227 | 86 | 6.9 ± 4.1 | 6 ± 0.6 |
| NA | A235 G10 1 | Single cell isolate after 10 consecutive batch fermentations with A-16235 | NA | ND | 93 ± 0.4 |
| NA | A235 G10 2 | Single cell isolate after 10 consecutive batch fermentations with A-16235 | NA | ND | 93 ± 0.1 |
| NA | A235 G10 3 | Single cell isolate after 10 consecutive batch fermentations with A-16235 | NA | ND | 83 ± 0.5 |

**Table 2.**
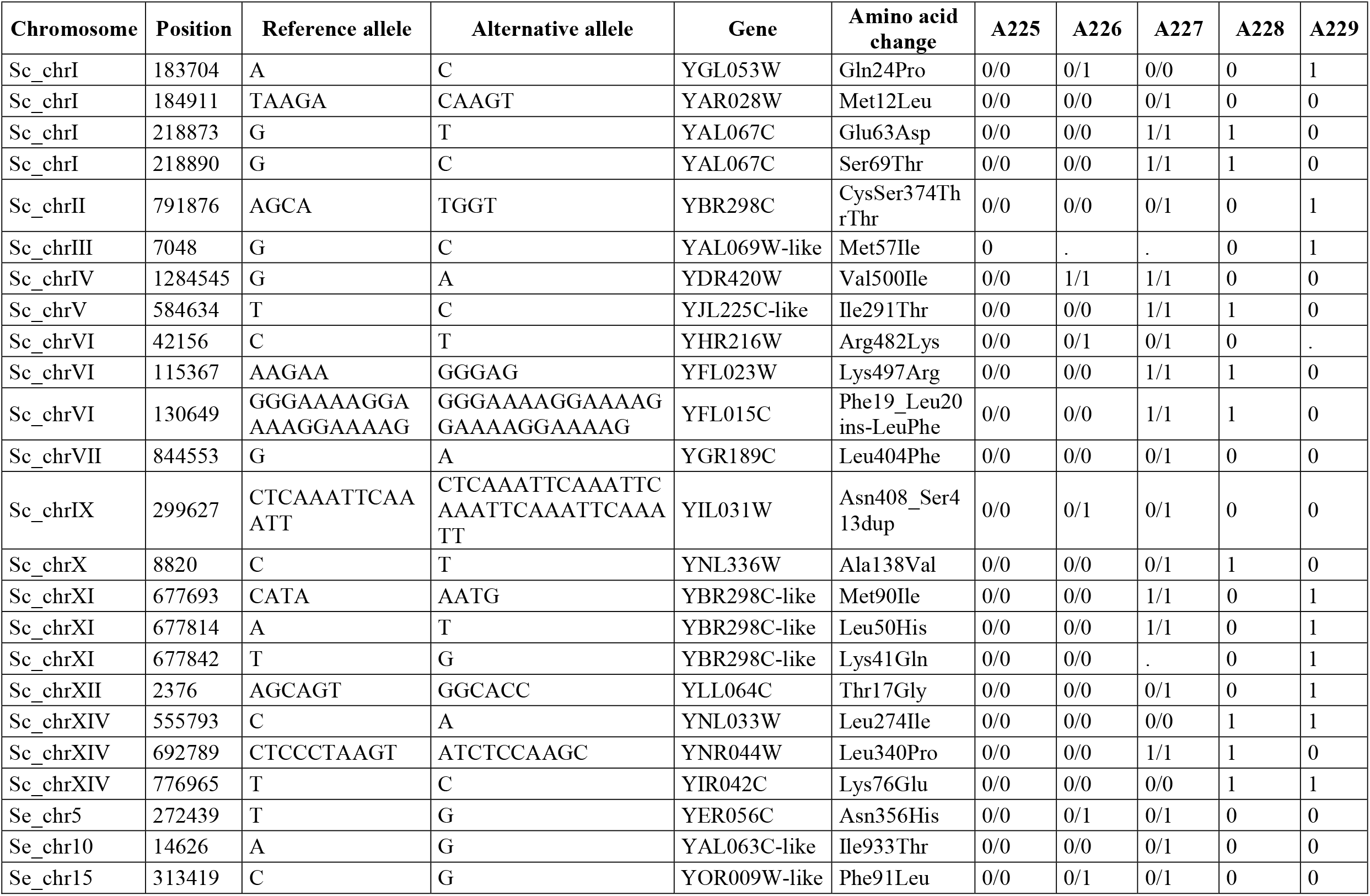
*de novo* SNPs in F1 spore clones of *S. cerevisiae × S. eubayanus* A225 hybrid.

A total of 1726 heterozygous SNPs were observed in the A227 F1 spore clone which was sporulated to produce the F2 spore clones A232-A235. However, a vast majority of these SNPs remained heterozygous in all four spore clones (1337), and only 38 segregated in a 2:2 pattern. In contrast to A227, only 8 *de novo* SNPs were detected in the four F2 spore clones. Of these, seven were intergenic and one a silent mutation. Hence, the four F2 spore clones were almost identical to A227 at a single nucleotide level, suggesting that any phenotypic differences between A227 and the four F2 spore clones are a result of larger-scale genomic variations.

Among the three single cell isolates of A235 that had undergone 10 consecutive batch fermentations in 15 °Plato wort, a total of 33 *de novo* SNPs were found. Only three of these SNPs were shared between all three single cell isolates. Of the 33 SNPs, three were missense mutations, one was a conservative inframe deletion, and one a conservative inframe insertion (Table 3). The affected genes include *PYC1* (YGL062W), encoding a pyruvate carboxylase. Of the remaining, twenty were intergenic and eight were silent mutations.

**Table 3.** *de novo* SNPs in G10 single cell isolates derived from the F2 spore clone A235.

| **Chromosome** | **Position** | **Reference allele** | **Alternative allele** | **Gene** | **Amino acid change** | **A235** | **A235 G10 1** | **A235 G10 2** | **A235 G10 3** |
| --- | --- | --- | --- | --- | --- | --- | --- | --- | --- |
| Sc_chrV | 584552 | CT | AC | YJL225C-like | p.Leu264Thr | 0/0 | 0/1 | 0/0 | 0/0 |
| Sc_chrV | 584565 | T | G | YJL225C-like | p.Val268Gly | 0/0 | 0/1 | 0/0 | 0/0 |
| Sc_chrVII | 386689 | TTGAT | TT | YGL062W | p.Asp672del | 0/0 | 0/0 | 0/0 | 1/1 |
| Sc_chrX | 8832 | G | A | YNL336W | p.Arg142Lys | 0/0 | 0/1 | 0/1 | 0/1 |
| Sc_chrXII | 1050334 | CTG | CTGTTG | YLR437C | p.Gln18dup | 0/0 | . | 1/1 | 0/0 |

Structural variations (SVs) in the *S. cerevisiae* A62 parent strain were estimated from the long reads using Sniffles. A total of 94 heterozygous SVs were identified, including 67 deletions, 27 insertions, 3 inversions, 1 duplication and 1 translocation (Supplementary Data 2). These SVs affected 18 genes, and the following cellular component GO terms were significantly enriched among the list: extracellular region (GO:0005576; *p*-value 1.2e-5), anchored component of membrane (GO:0031225; *p*-value 6.4e-4), fungal-type cell wall (GO:0009277; *p*-value 8.2e-4) and cell wall (GO:0005618; *p-*value 0.001). SVs in the F1 hybrid and derived spore clones were estimated from split and discordant Illumina reads using LUMPY through smoove. A total of 39 SVs were detected across the twelve strains (F1 hybrid, F1 spore clones, F2 spore clones, and G10 isolates), including 24 deletions, 2 duplications and 13 translocations (Supplementary Data 3). 12 deletion calls in the *S. cerevisiae* sub-genome of the F1 hybrid were supported by the SVs called for the A62 parent strain using the long reads. Of the 39 SVs in the hybrids, only five were absent from the F1 hybrid, suggesting few *de novo* SVs were formed during meiosis and the 10 consecutive batch fermentations in wort. While there was evidence of recombination within the *S. cerevisiae* sub-genome in the F1 and F2 hybrids, no recombination between the sub-genomes appears to have taken place, as indicated by the lack of split reads mapping to chromosomes from both sub-genomes.

In addition to the above mentioned SVs in the *S. cerevisiae* A62 parent strain, a heterozygous horizontal gene transfer event was observed on the right arm of chromosome XV, which contained an approx. 155 kbp region derived from *Torulaspora microellipsoides* (Supplementary Figure S3). This region includes the shorter 65 kb HGT region C that was originally described in *S. cerevisiae* EC1118 (Novo et al., 2009; Marsit et al., 2015) and is similar in size to the one later observed in *S. cerevisiae* CFC (a brewing strain) as a likely ancestral event (Peter et al., 2018). Because of heterozygosity, only two of the F1 spore clones (A226 and A229) carry this HGT region (Supplementary Figure S4). The presence of the HGT region C in wine yeast has been shown to improve oligopeptide utilization during wine fermentations (Marsit et al., 2015), yielding an advantage in nitrogen-limited media, but its effect in wort fermentations remains unclear.

### Phenotypic variation in the strain breeding panel

A range of brewing-relevant industrial phenotypes were assessed in the twelve *de novo* hybrids and the parent strains. These 22 phenotypes included consumption and uptake of maltose and maltotriose, fermentation rate, flocculation, viability, growth at 4 and 37 °C, and formation of eleven aroma-active compounds. Extensive phenotypic variation was observed between the strains (Figure 4). Both hierarchical clustering based on Euclidean distance (Figure 4A) and principal component analysis (Figure 4B-C) grouped the F1 hybrid in between the parent strains, while F1 and F2 spore clones grouped around the strain they were derived from (A225 and A227, respectively). As has been observed in previous studies on *de novo* brewing yeast hybrids (Mertens et al., 2015; Krogerus et al., 2016, 2018b), both mid-parent and best-parent heterosis was observed among the different hybrid strains and the various phenotypes.

**Figure 4.**
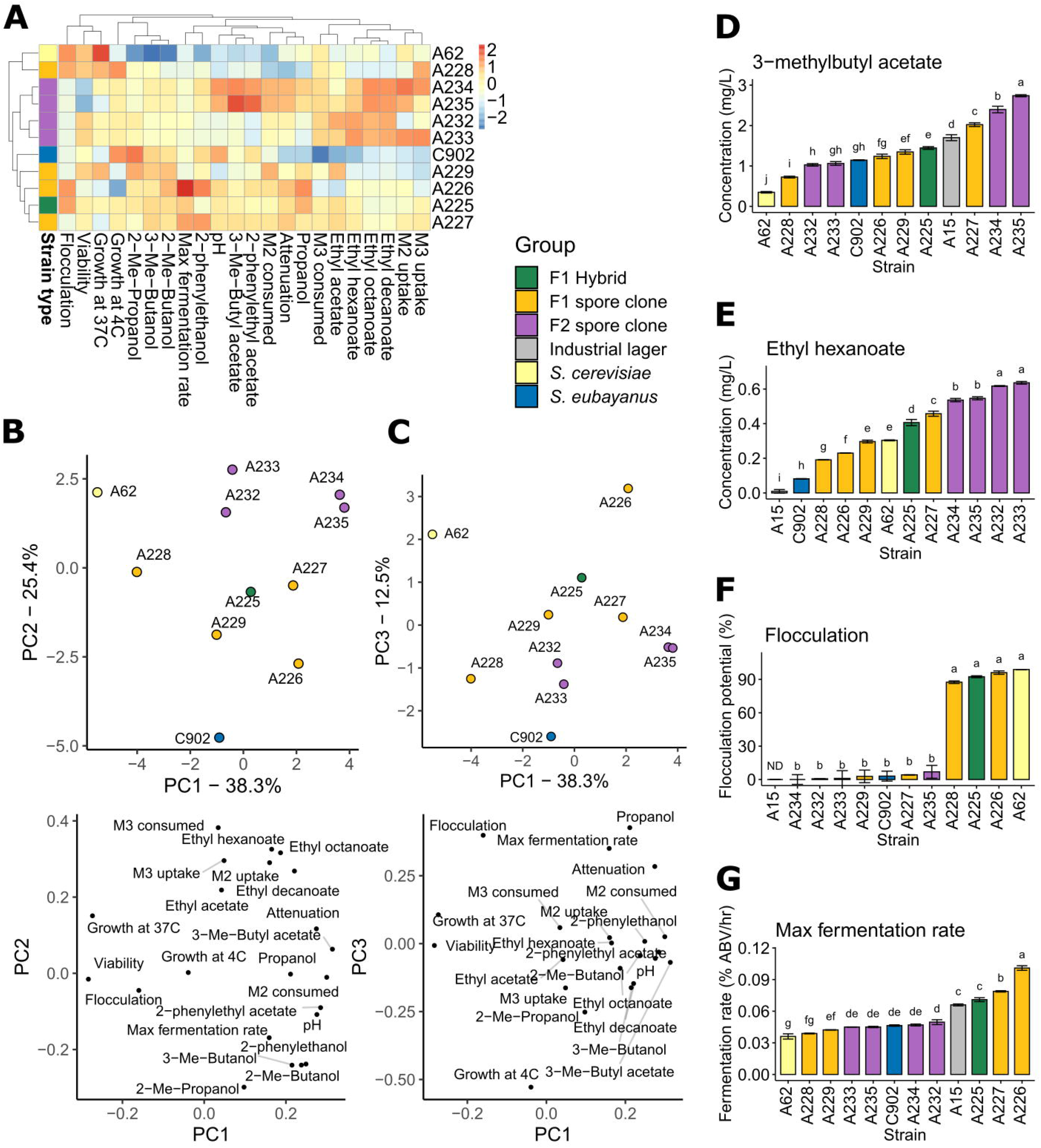
Phenotypic variation in the parent strains and hybrids. (A) Heatmap depicting the variation of the 22 phenotypic traits in the parent strains, F1 hybrid, F1 spore clones and F2 spore clones. (B and C) Principal component analysis of the 22 phenotypic traits. (D) 3-methylbutyl acetate and (E) ethyl hexanoate concentrations in the beers produced with the above 11 strains and a commercial lager yeast control. (F) The flocculation potential of the above 11 strains as measured by Helm’s test. (G) The maximum fermentation rate observed among the above 11 strains and a commercial lager yeast control during the wort fermentations. (D-G) Values are means from two independent fermentations and error bars where visible represent the standard deviation. Values with different letters (a–j) above the bars differ significantly (*p* < 0.05) as determined by one-way ANOVA and Tukey’s test.

#### Aroma diversity

Interest towards beer with novel and diverse flavours is increasing (Aquilani et al., 2015; Carbone and Quici, 2020; Gonzalez Viejo and Fuentes, 2020), and the results here suggest that hybridization and subsequent sporulation can give rise to lager yeast strains with both enhanced and diverse production of aroma-active compounds. 3-methylbutyl acetate, with its banana- and pear-like aroma, is one of the most important yeast-derived flavor compounds in beer (Pires et al., 2014). Here, we measured higher concentrations of this ester in the beer produced with the F1 hybrid A225 compared to either of the parent strains (Figure 4D). Of the four F1 spore clones, one (A227) produced higher levels of 3-methylbutyl acetate than the F1 hybrid. The F1 strain A227 was chosen for further sporulation and spore clone screening due to its high production of 3-methylbutyl acetate. Two out of four F2 spore clones produced the highest levels of 3-methylbutyl acetate among all tested strains, reaching 2.5-fold higher levels than the most productive parent strain (*S. eubayanus* C902). This ester was produced only at very low levels by the *S. cerevisiae* A62 parent strain.

Similarly to 3-methylbutyl acetate, considerable variation was observed for ethyl hexanoate formation. Ethyl hexanoate, with its apple- and aniseed-like aroma, is another important yeast-derived flavour compound in beer (Pires et al., 2014). Again, the F1 hybrid produced higher concentrations of this ester compared to either parent strain (Figure 4E). Of the F1 spore clones, A227 again produced the highest levels of ethyl hexanoate, while the highest levels among all tested strains was observed in the four F2 spore clones derived from A227. Two-fold higher ethyl hexanoate levels were observed in the beers made from these strains compared to the better parent strain (*S. cerevisiae* A62). Low concentrations of this ester were produced by the *S. eubayanus* C902 parent strain and the industrial control *S. pastorianus* A15.

As 3-methylbutyl acetate and ethyl hexanoate formation was strongly associated with the two parent strains, *S. eubayanus* C902 and *S. cerevisiae* A62, respectively, hybridization yielded a strain producing high levels of both. Interestingly, a strain producing several-fold higher levels of both these esters could be derived by selecting meiotic segregants. Highest concentrations of ethyl hexanoate were seen with the four F2 hybrids. In the case of 3-methylbutyl acetate, the highest concentrations were also seen in F2 hybrids, though in this case only for the two tetraploid strains.

#### Fermentation performance

In addition to greater aroma diversity, brewers also demand strains with efficient fermentation. As expected based on previous studies with similar hybrids (Krogerus et al. 2015, 2016, 2017), the tetraploid strain A225 fermented wort more rapidly and completely than the parental strains (Figure 4A and 4G). Alcohol level at the end of the hybrid fermentation was 6.7% (v/v) compared to 5.7% and 4.9% for the ale and *S. eubayanus* strain respectively. A direct comparison of the fermentation performance of the tetraploid hybrid and four F1 sibling strains revealed clear differences that were associated with ploidy. The maximum fermentation rate of the tetraploid F1 siblings was slightly higher than that of the parental hybrid (Figure 4G). Alcohol level was higher relative to the parent (approx. 6.5% compared to 6.2%). Fermentation rates of the diploid strains were similar to that of the parental tetraploid in the early stage of the fermentation (up to 72h), but were lower thereafter. Final yields of alcohol in the strains A228 and A229 were 4.2 and 4.4%, respectively. Similarly to the F1 spore clones, the fermentation performance of the F2 spore clones appeared to be associated with ploidy. While little difference was seen in the maximum fermentation rates (Figure 4G), due to similar performance early in fermentation, the tetraploid strains A234 and A235 finished at higher alcohol levels (7.0 and 6.9%, respectively) compared to the diploid strains A232 and A233 (6.0 and 5.7%, respectively). Of the *de novo* hybrid strains, A225-A227 all outperformed the industrial lager yeast A15 that was included as a reference with respect to maximum fermentation rate.

#### Flocculation

The *S. cerevisiae* A62 parent showed strong flocculation, while flocculation potential was low in the *S. eubayanus* C902 parent strain. The F1 hybrid also showed comparably strong flocculation relative to the parent strain, and interestingly two out of the four F1 siblings showed strong flocculation, while the others showed weak flocculation (Figure 4F). Flocculation potential was not linked to the ploidy of the spore clones, suggesting that the heterozygous genotype of the *S. cerevisiae* A62 parent may be responsible. Indeed, a number of heterozygous SVs linked with extracellular region and cell wall were identified, including a 135 bp deletion in *FLO5* and a 65 bp deletion in *TIR2* (Supplementary Data 2), which could potentially explain this loss of flocculation in half the spore clones. A227 and the F2 spore clones and derived G10 isolates all exhibited weak flocculation. The *TIR2* deletion was identified from the short-read data, and was present in spore clones A226 (strong flocculation) and A227 (weak flocculation), however the *FLO5* deletion was not detected.

#### Spore viability

Both the domesticated strains studied here had a low level of sporulation and spore viability. In the A15 lager strain, sporulation was not observed and in the *S. cerevisiae* A62 ale strain, it was only observed at a low level (21%) and of these only 8% were found to be viable. In contrast, the sporulation efficiency of the *S. eubayanus* strain was high and spores were generally viable (Table 1). Sporulation in the A225 tetraploid strain was intermediate between the parents with spore viability measured as 55%. In the F1 and F2 generation, sporulation and spore viability was largely influenced by ploidy with spore viability ranging from 0% to 95%. Diploid strains were found to have low sporulation efficiency and to be sterile. An exception was the diploid F2 spore clone A232, which had a spore viability of 78% (Table 1).

### Phenotypic stability of an F2 spore clone

The phenotypic stability of the three G10 isolates of the F2 segregant A235, isolated after 10 consecutive fermentations in industry-strength all-malt wort, was assessed by comparing the isolates and the G10 mixed population to A235. In wort fermentations, the G10 mixed population did not perform as well as the original A235 strain, despite a relatively rapid fermentation rate in the first 72 hours (Figure 5A). The final alcohol yield was 6.9%, compared to 7.1% for the original strain. It was however clear that the G10 population was phenotypically heterogenous in nature. The three single cell isolates derived from the G10 population showed clearly different capacities to ferment the wort. Weakest performance was observed with isolate 2, best performance with isolate 3 and an intermediate performance with isolate 1. Aroma formation was also affected by the repeated wort fermentations. Significantly lower amounts of 3-methylbutyl acetate were formed by the G10 population and single cell isolates compared to A235 (Figure 5B), while ethyl hexanoate levels in the G10 isolates were similar or slightly lower than A235 (Figure 5C). Futhermore, while A235 was able to sporulate, none of the three single cell isolates produced ascospores when inoculated onto potassium acetate agar (Table 1).

**Figure 5.**
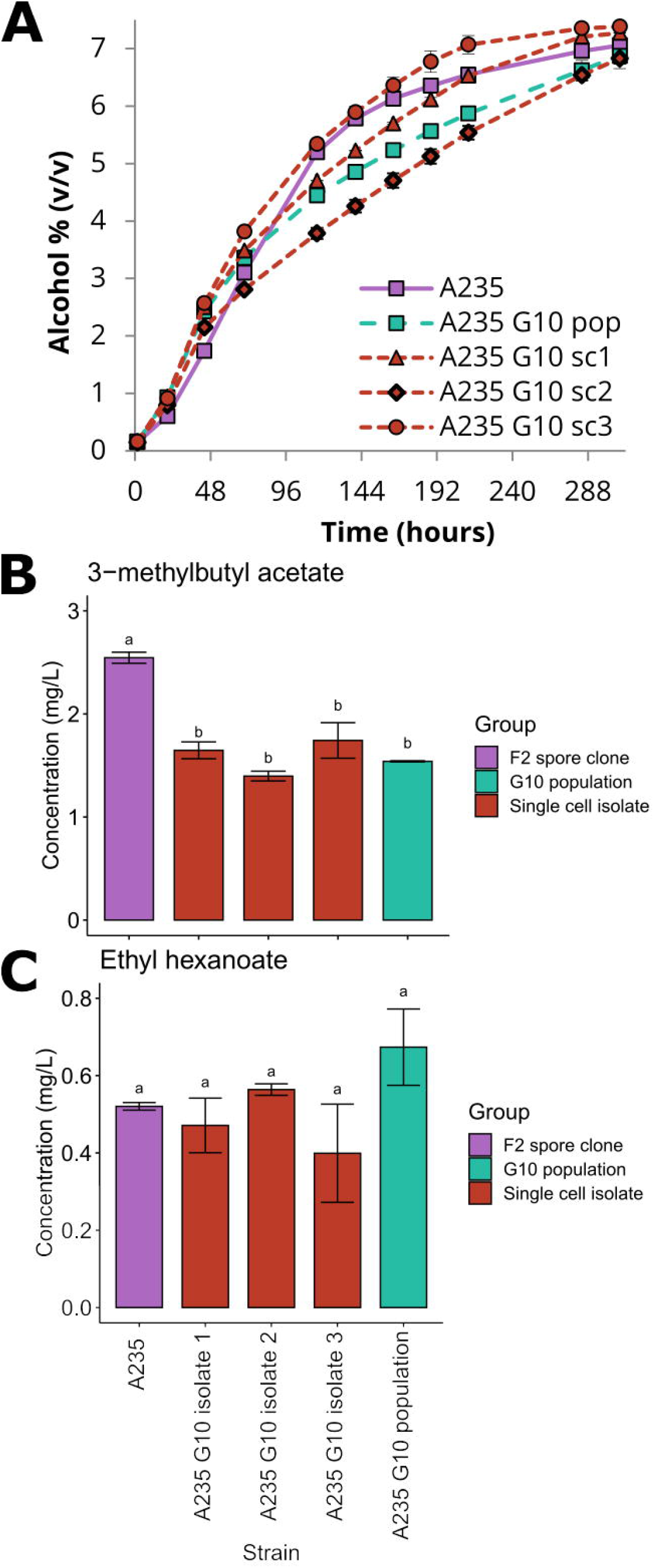
Fermentation performance of the G10 isolates and the mixed population. (A) The alcohol content (% volume) of the 15 °P wort fermented with the F2 spore clone A235, the tenth generation mixed population derived from it, and the three single cell isolates from the tenth generation population. (B) The 3-methylbutyl acetate and (C) ethyl hexanoate concentrations in the beers produced with the above strains. Values are means from two independent fermentations and error bars where visible represent the standard deviation. Values with different letters (a–b) above the bars differ significantly (*p* < 0.05) as determined by one-way ANOVA and Tukey’s test.

## Discussion

Limited phenotypic and genetic diversity exists between industrial lager yeasts (Okuno et al., 2015; Gallone et al., 2019; Langdon et al., 2019). In this study, we sought to explore how the fertility of a newly created tetraploid *S. cerevisiae × S. eubayanus* interspecies hybrid could be exploited to expand the phenotypic diversity of this group. Rare mating was used to produce a polyploid hybrid. This can occur, e.g. by inactivation of one *MAT* locus or through spontaneous gene conversion to produce parental strains that are homozygous for mating type (*MAT*a*/MAT*a or *MATα/MATα*) (Gunge and Nakatomi, 1972; Greig et al., 2002; Sipiczki, 2018). In the current study, rare mating appears to have been facilitated through the former mechanism. Sequencing of the F1 hybrid suggests that one *MAT* locus in the diploid parental *S. cerevisiae* cell was lost through whole-chromosome deletion of chromosome III, effectively producing a cell that was hemizygous for mating type. Similar losses of the same chromosome have also recently been observed in artificial *S. cerevisiae* × *S. kudriavzevii* and *Saccharomyces kudriavzevii x Saccharomyces uvarum* hybrids (Karanyicz et al., 2017; Morard et al., 2020). What induced the parental *S. eubayanus* cell to engage in rare mating remains unclear. Loss of one copy of chromosome III has previously been observed in allotriploid and allotetraploid hybrids derived from the A62 ale strain (Krogerus et al., 2016). The strain, therefore, appears susceptible to this change and, as a result, is particularly suitable for natural allopolyploid hybridization. To what extent chromosome III loss is responsible for hybridization in interspecies hybrids requires further investigation.

As observed in previous studies on allotetraploid yeast (Greig et al., 2002; Sebastiani et al., 2002; Antunovics et al., 2005; Naseeb et al., 2021) there appeared to be no post-zygotic barrier to reproduction with the F1 hybrid investigated here. Fertility of the F1 spore clones was also limited to tetraploid strains (via endomitosis (Sebastiani et al., 2002) or, as is most likely the case here, self-fertilization of homo- or hemizygous diploid spores). Interestingly, fertile strains were observed among both diploid and tetraploid F2 spore clones. Antunovics et al. (2005) showed persistent fertility of a presumed alloploid hybrid over several generations, though in that case the fertility was restricted to allotetraploid cells. The mechanisms that facilitate this phenomenon are not yet known but appear to be unrelated to chromosome pairing as fertility was not directly influenced by ploidy (Greig et al. 2002). Further investigation is necessary to elucidate the processes involved, and may even help to clarify those processes that contribute to speciation. Marcet-Houben & Gabaldón (2015) have, for example, suggested that an ancient interspecies hybridization may have led to the creation of the ancestral *S. cerevisiae* lineage. Regardless of the mechanisms involved, generation of allotetraploid hybrids appears to be potentially useful for generating diversity through meiotic recombination (Bozdag et al., 2021; Naseeb et al., 2021). Here, no evidence of recombination between the two parental sub-genomes of the hybrid was observed, rather only within the parental sub-genomes.

Industrial lager beer fermentation is currently dominated by Frohberg-type *S. pastorianus* strains, and there exists little diversity within the group (Gallone et al., 2019; Langdon et al., 2019). Creating new flavour profiles, e.g. in response to the increased consumer demand for higher product quality and beer with novel and diverse flavours (Aquilani et al., 2015; Carbone and Quici, 2020; Gonzalez Viejo and Fuentes, 2020), is hampered by the low level of diversity amongst commercial brewing yeast strains. Previous research has shown that interspecific hybridization is an effective way of introducing new aromatic diversity among lager yeasts (Krogerus et al., 2015; Mertens et al., 2015; Nikulin et al., 2018; Turgeon et al., 2021). Not only can distinct aroma profiles of different parent strains be combined, but aroma formation is often improved compared to either of the parents from heterosis. Here, we show that sporulation of fertile allotetraploid hybrids could be exploited to further improve aroma production, as beer concentrations of two important aroma-active esters 3-methylbutyl acetate and ethyl hexanoate were up to 2.5-fold higher in the F2 spore clones compared to the best parent. The variation between spore clones can also be exploited to tailor the *de novo* hybrid towards specific desired traits. It must, however, be emphasised, that much of the phenotypic variation observed here was likely due to segregation and loss-of-heterozygosity in the heterozygous *S. cerevisiae* sub-genome.

Phenotypic stability is an essential trait in any industrial yeast and this is particularly relevant for interspecies hybrids where genomes are known to be inherently unstable. Here, the stability of the F2 spore clone A235 was assessed after consecutive wort fermentations. The results showed clearly differences in performance between A235 and the G10 population but also between the single-cell cultures. Differences were evident for fermentation capacity, flocculation and flavour profile and were not due to structural variation as no such changes were apparent. There were however several CNV changes with respect to chromosomes. The single-cell cultures all gained two extra copies of *S. eubayanus* chromosome III. Isolate 1 lost both copies of the *S. cerevisiae* chromosome XII, while Isolate 2 lost two copies of *S. eubayanus* chromosome XII. Morard et al. (2019) also observed that copy number gains of chromosome III resulted in increased ethanol tolerance, possibly from upregulation of stress-related genes located on it. Voordeckers et al. (2015) in a study of ethanol adaptation also noted changes in the number of these same chromosomes. In response to high ethanol, several strains independently gained copies of one or both of these chromosomes. The authors suggested that these changes may be an early adaptive response to ethanol, which would be followed by more refined changes with additional exposure. It may be that the G10 yeast in this study are similarly showing signs of early adaptation to ethanol, which reached up to and over 7% in these fermentations. The higher cell viability of G10 populations is consistent with an improved tolerance, though the exact relationship between these specific CNVs and phenotype has yet to be resolved.

Genomic stability of brewing yeast is vital from an industrial point-of-view. This is because, in contrast to other beverage fermentations, brewing yeast is reused for multiple consecutive fermentations. The instability that was demonstrated here for the tetraploid F2 segregant A235, highlights the importance of stabilizing *de novo* yeast hybrids before they are suitable for industrial use. While instability is not a desirable trait for industrial yeast, rapid genome resolution in interspecies hybrids, such as that seen in this and other studies (Dunn et al., 2013; Peris et al., 2017; Smukowski Heil et al., 2017), suggests that stable genomes may evolve within a short time and, furthermore, that *de novo* hybrid genomes may be amenable to directed evolution to improve their industrial potential (Krogerus et al., 2018a; Gorter de Vries et al., 2019). This opens up the possibility of further improving and developing the strains in a targeted manner.

A key feature of the modern brewing market is a demand for diversity in beer character. Until now brewers have satisfied this demand through the creative use of malts and hops. This study, and related investigations, have shown that there is also significant potential to direct or fine-tune the flavour profile of beers through the creation of novel brewing yeast strains or modification of existing brewing yeast strains. Here, a number of development steps were undertaken (hybridization, sporulation, adaptation) to introduce diversity. It is clear however that further improvement may be achieved through the addition of even more developmental steps, e.g. further rounds of sporulation, or evolutionary engineering. Importantly, all stages in the strain development included here could feasibly occur in nature. Strains thus produced are therefore suitable for immediate application in brewing, with the proviso that genome stabilization has occurred prior to application. Further investigation is required to determine the dynamics of genome stabilization following hybridization.

## Acknowledgements

Eero Mattila and Niklas Fred are thanked for assistance in the VTT Pilot Brewery, and Aila Siltala for skilled technical assistance.

## Author Contributions

Conceived the study: BG

Designed experiments: KK, BG

Performed experiments: KK, FM, VV, BG

Analysis of experimental data: KK

Analysis of genome data: KK, SC, GP, MDC, JXY, GL

Wrote the manuscript: KK, BG

