## Supplemental Tables and Figures for "Lager yeast design through meiotic segregation of a fertile *Saccharomyces cerevisiae* x *Saccharomyces eubayanus* hybrid"

Keywords:

**Supplementary Table S1**. Assembly statistics for *S. cerevisiae* A81062

| | |
|---|---|
| Genome size (bp) | 12646885 |
| Contigs | 21 |
| Mean (bp) | 602232 |
| Median (bp) | 586058 |
| N50 (bp) | 917785 |
| Largest (bp) | 1456411 |
| GC(%) | 38.15 |

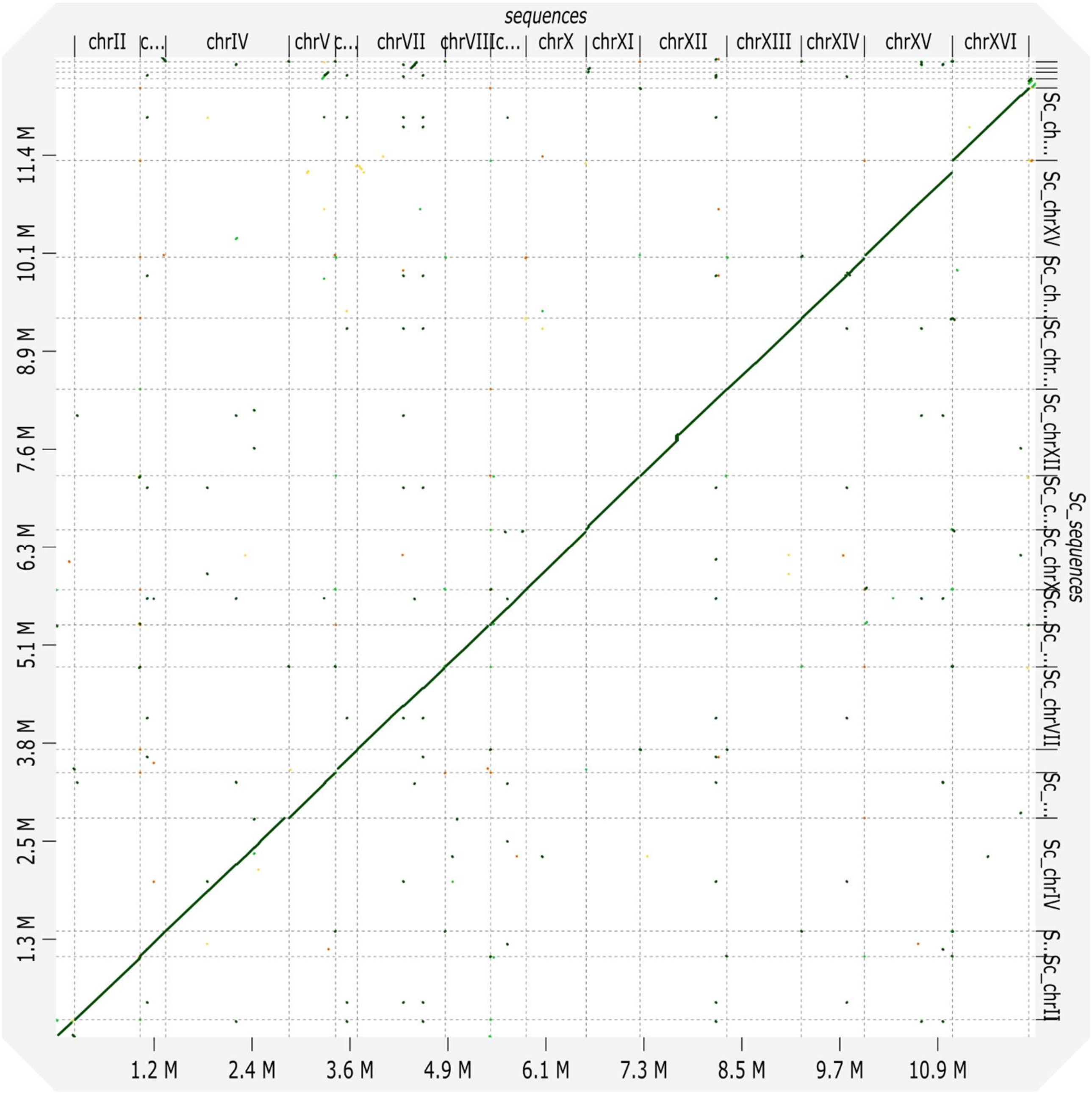


**Supplementary Figure S1** - Alignment of *S. cerevisiae* A81062 assembly contigs (y-axis) to *S. cerevisiae* S288C (x-axis)

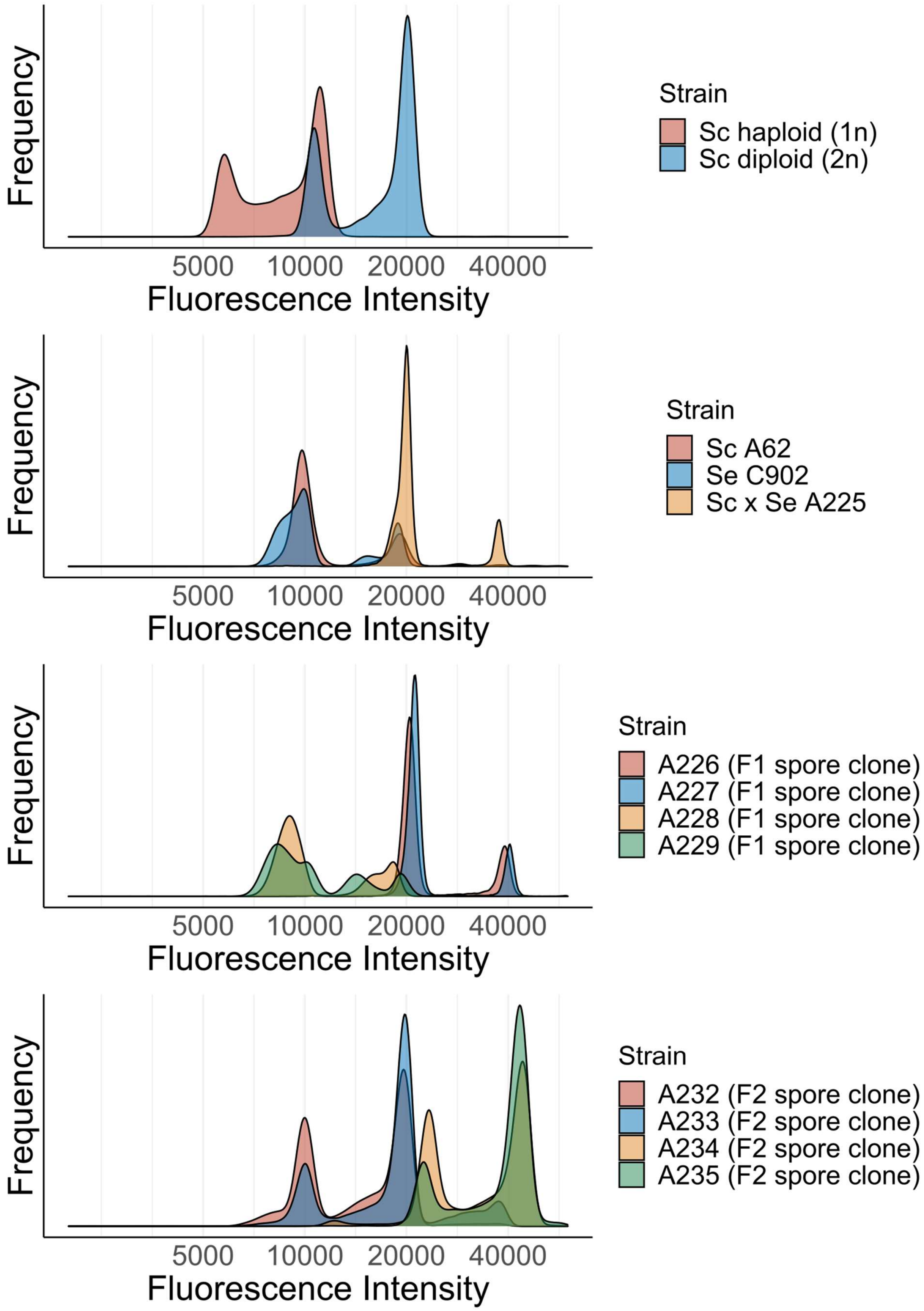


**Supplementary Figure S2** - Fluorescence intensity after SYTOX Green-staining

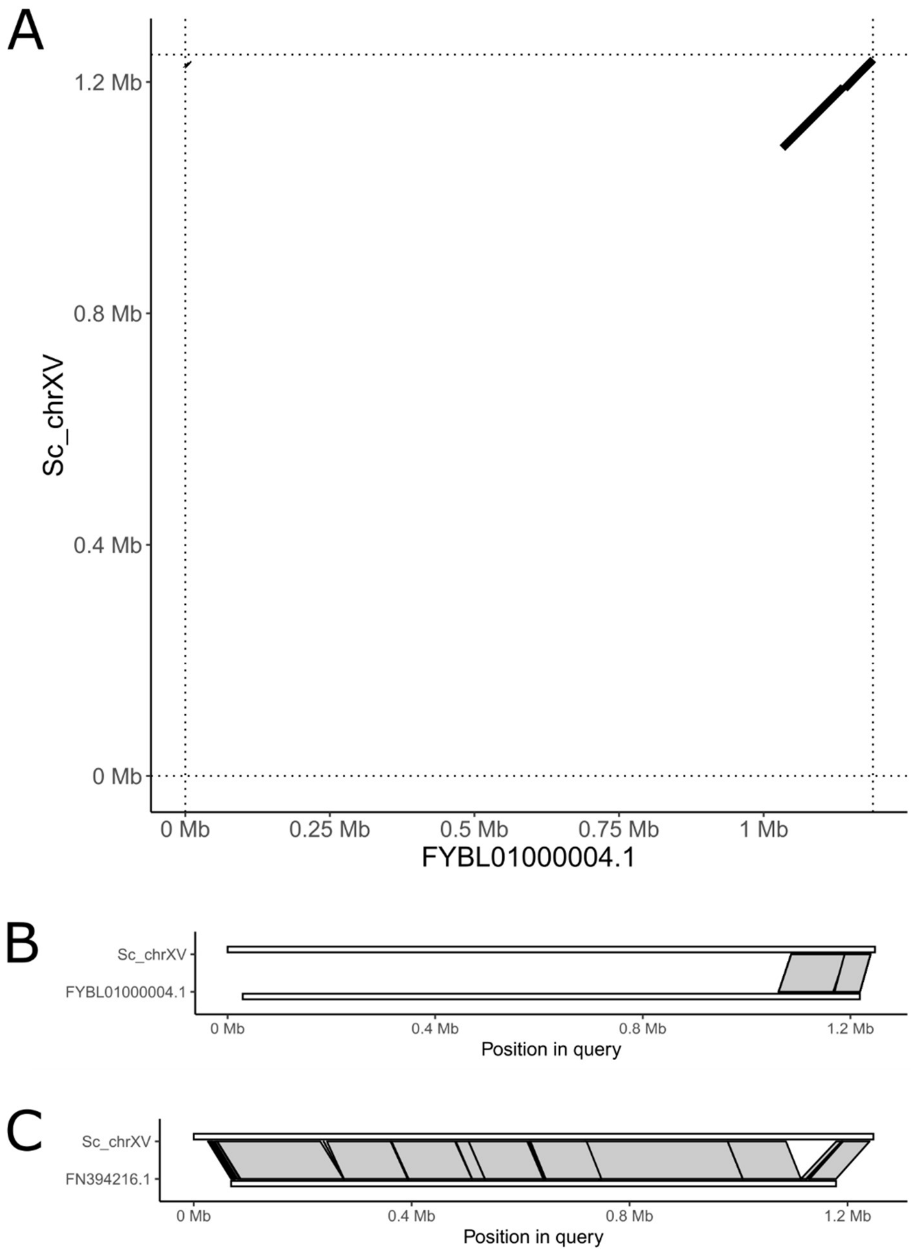


**Supplementary Figure S3** - (A and B) Alignment of chrXV of *S. cerevisiae* A81062 to *Torulaspora microellipsoides* scaffold 2 (FYBL01000004.1). (C) Alignment of chrXV of *S. cerevisiae* A81062 to *S. cerevisiae* EC1118 scaffold (FN394216.1)

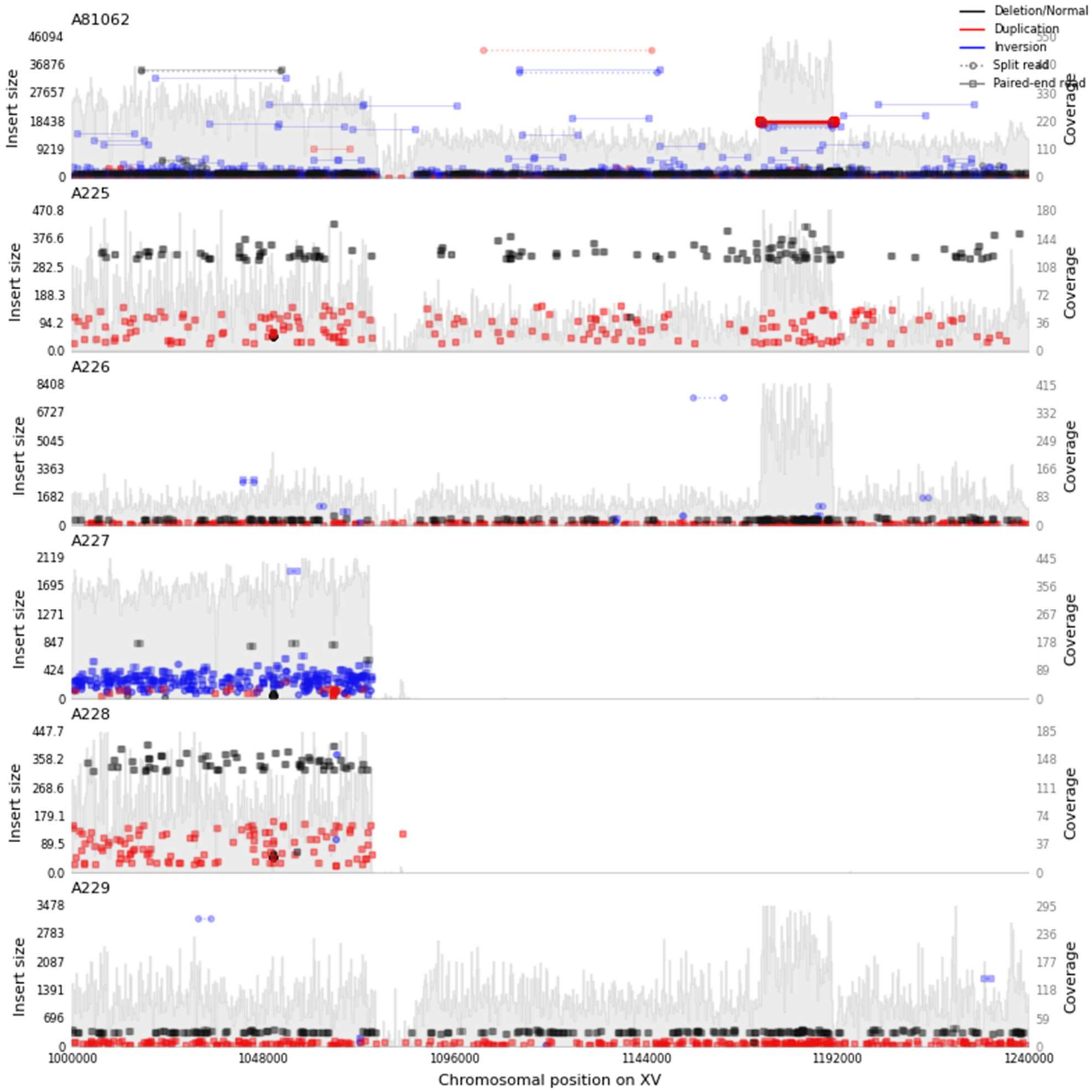


**Supplementary Figure S4** - Sequencing coverage on the right arm of *S. cerevisiae* A81062 chrXV in the F1 hybrid A225 and derived F1 spore clones.
